# Vector dynamics influence spatially imperfect genetic interventions against disease

**DOI:** 10.1101/2020.07.16.207464

**Authors:** Mete K Yuksel, Christopher H Remien, Bandita Karki, James J Bull, Stephen M Krone

**Affiliations:** Department of Mathematics, University of Idaho, Moscow, ID 83844-1103 USA; Department of Biological Sciences, University of Idaho, Moscow, ID 83844-1103 USA

**Keywords:** genetic pest management, gene drive, pathogen suppression, mosquito biting dynamics, spatial structure, Ross-Macdonald, mathematical model

## Abstract

**Background and objectives:** Genetic engineering and similar technologies offer promising new approaches to controlling human diseases by blocking transmission from vectors. However, in spatially structured populations, imperfect coverage of the vector will leave pockets in which the parasite can persist. Yet movement by humans may disrupt this local persistence and facilitate eradication when these pockets are small, essentially distributing parasite reproduction out of unprotected areas and into areas that block its reproduction.

**Methodology:** We develop formal mathematical models of this process similar to standard Ross-Macdonald models, but (i) specifying spatial structure of two patches, with transmission blocked in one patch but not in the other, (ii) allowing temporary human movement (travel instead of migration), and (iii) considering two different modes of mosquito biting.

**Results:** We find that there is no invariant effect of disrupting spatial structure with travel. For both biting models, travel out of the unprotected patch has different consequences than travel by visitors into the patch, but the effects are reversed between the two biting models.

**Conclusions and implications:** Overall, the effect of human travel on the maintenance of vector-borne diseases in structured habitats must be considered in light of the actual biology of mosquito abundances and biting dynamics.

**Lay summary:** Genetic interventions against pathogens transmitted by insect vectors are promising methods of controlling infectious diseases. These interventions may be imperfect, leaving pockets where the parasite persists. How will human movement between protected and unprotected areas affect persistence? Mathematical models developed here show that the answer is ecology-dependent, depending on vector biting behavior.

## Introduction

Radically new technologies are becoming available to suppress vectored diseases. They operate as genetic modifications of vector populations that block parasite transmission. One such technology uses ‘modification’ gene drives that automatically sweep through the population. The drive is engineered to include one or more genes that interfere with the parasite in the vector (Gould, 2008; Burt, 2014; Gantz et al., 2015). A somewhat parallel approach, but without genetic engineering, introduces pathogen-blocking strains of the self-spreading bacterial symbiont *Wolbachia* into the vector (Hoffmann et al., 2011; Schmidt et al., 2017). A third, and more mundane approach is to release huge numbers of lab-reared, genetically modified vectors, simply to infuse wild populations with transmission-blocking genes in a manner akin to the sterile insect technique (Evans et al., 2019; Gould et al., 2006). The gene drive and *Wolbachia* approaches result in possibly permanent alterations of vector populations because the genetic modifications are selectively maintained. The swamping method is typically transient, because the modification is not coupled with any selective benefit (Gould et al., 2006); continual releases of engineered vectors would be required to maintain the parasite block.

Genetic modifications have an advantage in that they accrue directly and specifically to the vector and are transmitted intact to offspring, contrasting with pesticides that are broadcast environmentally, cannot be uniformly applied and need to be applied repeatedly. However, genetic methods are sometimes controversial and face extreme regulatory hurdles because of their transgenerational permanence. We nonetheless imagine that many of these genetic technologies will be widely implemented in the near future, so predicting the possible bases of failure versus success may be useful in ensuring the best possible outcomes. Some methods may seem so foolproof as to ensure disease eradication because of their ability to modify huge fractions of vector populations. Even so, one worry is that any population intervention is likely to be somewhat incomplete, leaving spatial pockets of minimal coverage interspersed with perhaps large pockets of almost total coverage (e.g., North et al., 2013, 2019). What will be the effect of these pockets of poor coverage? From a greatly simplified spatial model of pathogen dynamics, we previously suggested that spatial structure will foster the persistence of the pathogen when the pathogen would disappear in the absence of structure (Bull et al., 2019). Thus, any softening of spatial structure would help limit parasite persistence. That model omitted vectors as well as hosts, so any inference to vector dynamics was tangential. Here we consider a more realistic model of spatial structure than we addressed previously: a model that includes vectors, with host mobility; when hosts are spatially clustered and a genetic intervention blocks vector transmission most places, does host movement invariably facilitate eradication? Furthermore, how does the effect of human movement depend on the transmission dynamics?

Our question has many precedents in previous mathematical models of vectored diseases, of which the Ross-Macdonald models are the original and most prominent (Keeling and Rohani, 2008). The effect of spatial structure on disease dynamics has been addressed in several modeling studies when assuming a single model of transmission dynamics (Cosner et al., 2009; Prosper et al., 2012; Ruktanonchai et al., 2016; Anzo-Hernández et al., 2019; Soriano-Paños et al., 2020; Khamis et al., 2020). The effect of different models of transmission dynamics has been addressed in the absence of spatial structure (McCallum et al., 2001; Wonham et al., 2006). Our models will combine spatial structure, differential blocking of transmission among patches, human movement among patches, and different forms of mosquito biting dynamics. Our assemblage of assumptions is unique, but this broad foundation of previous work simplifies our task and provides many anchor points to validate our findings.

## Results

### Foundations

The Introduction provided several biological contexts for the problem we study. They all involve vectored infectious diseases, spatial structure, and movement of vectors and/or humans (we consider only the latter here). Here we explain how that biology is converted into our models.

### Population structure

Our models are standard epidemiological ‘SIS’ models, accounting for vector (mosquito) and host (human) numbers, as well as spatial structure. Parasites have no individual existence *per se* in the model; they exist only as infected states of mosquitoes or humans. Infections are transmitted only mosquito to human or human to mosquito. A full description of the mathematical models is given in the Appendix.

To abstract this biological process, we model a population with discrete subpopulations; the same population subdivisions coincide for both humans and vectors, but it operates somewhat differently for humans than for vectors. The number of humans in each patch is invariant; no one is born and no one dies during the time period considered. In contrast, mosquitoes have a patch-specific birth rate (independent of the number of mosquitoes and humans) and a patch-invariant death rate, leading to a patch-specific equilibrium density; mosquito lifetimes, on the order of weeks or months, are much shorter than human lifetimes. Mosquito spatial structure is rigid and invariant, whereas humans have a home patch but travel temporarily to non-resident locations—a movement scheme that differs from formal ‘migration’ (Cosner et al., 2009). The state of mosquito infections at a location depends on mosquito behavior and on the history of their exposure to humans at that location, regardless of whether the humans were residents or visitors. In contrast, humans are not confined to one location throughout life; they move, but each person is identified with a home residence, regardless of their location at any moment. This process would arise with daily commuting, jobs that involve travel, and even some kinds of nomadic lifestyles. (Our approach thus differs from standard migration models in which individuals move without memory of an individual’s previous residence.) Because humans travel, their infection status depends on their history of exposure to mosquitoes at the different locations they have occupied.

### Transmission dynamics

We consider two models of infection dynamics as they affect mosquito biting rates: density-dependent and frequency-dependent (McCallum et al., 2001; Wonham et al., 2006). These models differ in the way the biting rate of mosquitoes at a site scales with the number of humans at that site (Fig. 1). In the density-dependent model, characterized by a mass-action functional response, the rate at which a single person is bitten is independent of the number of humans; in the frequency-dependent model, characterized by a saturated functional response, the total number of bites is determined by the number of mosquitoes, so adding more humans decreases the bite rate per person unless mosquito density increases with human density. The standard Ross-Macdonald models often assume frequency-dependence.

**Figure 1:**
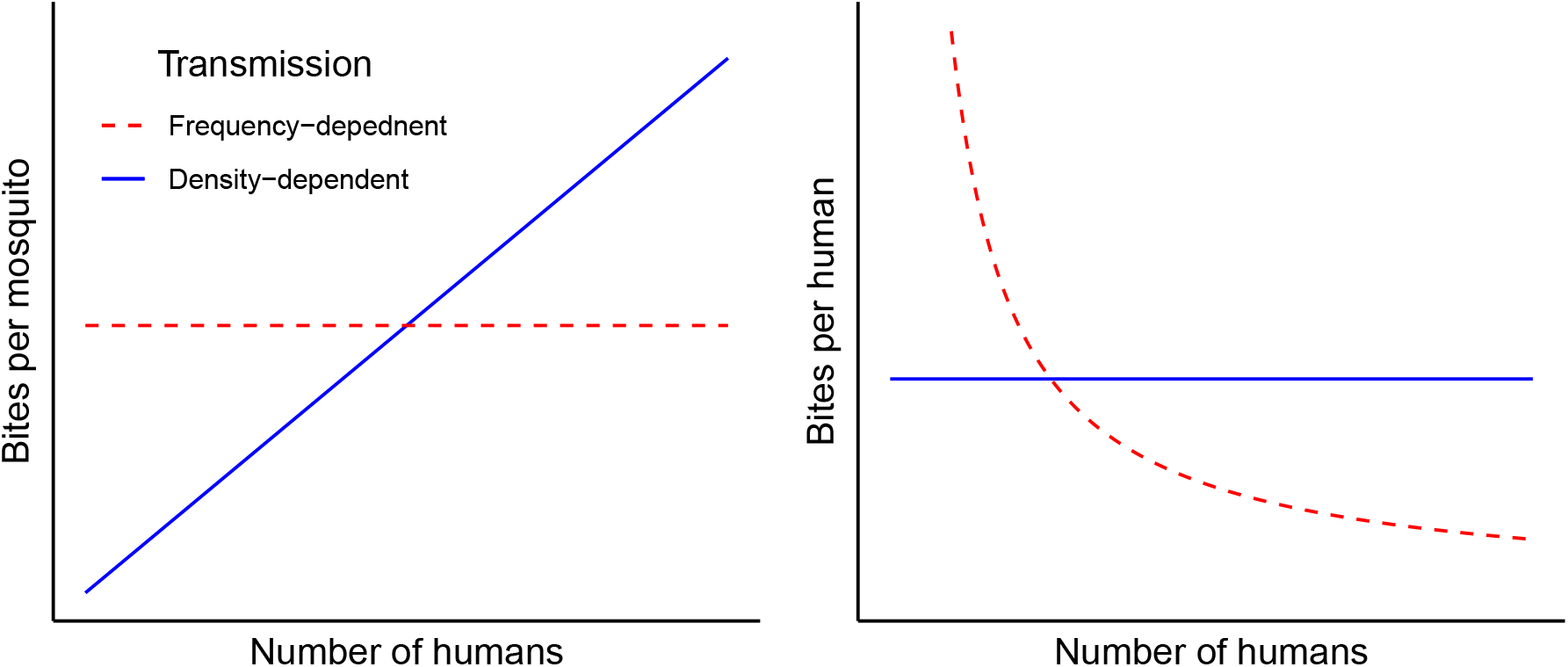
Differences between the frequency-dependent and density-dependent models with respect to biting dynamics. The left panel shows biting rate per mosquito, the right panel shows biting rate per human. The solid (blue) lines apply to the density-dependent case, dashed (red) to the frequency-dependent case.

**Figure 2:**
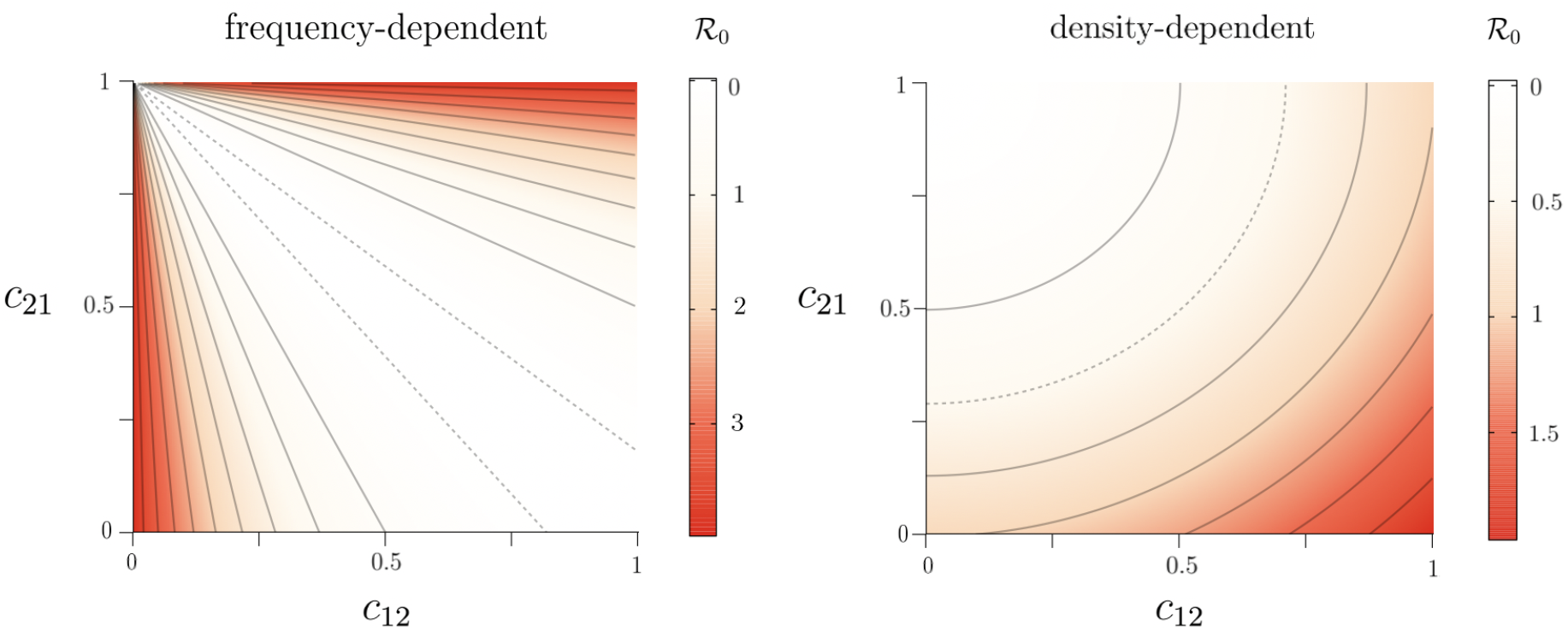
Contour plots of the basic reproduction number as a function of the visitation parameters (*c*_12_ and *c*_21_) reveal a fundamentally different effect of human movement under frequency dependence (left) than under density dependence (right). In each panel, the dashed contour line represents 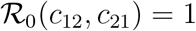, solid curves represent other values. These plots used a single set of parameters except for the *c*_*ij*_, but plots using other values are similar, except the curvature (and steepness in the FD case) of the contour lines change when other parameters are varied. Values of the parameters used are *γ* = 0.071, *λ*_1_ = 50 *λ*_2_ = 50, 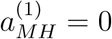, 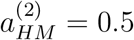, *a_HM_* = 0.8, *δ* = 0.02, *b_DD_* = 0.000075, and *b_FD_* = 0.0375. The population sizes used are *H*^(1)^ = *H*^(2)^ = 500 and *M* ^(2)^ = 2500. Calculations were done using expressions (1) and (2) for the basic reproduction numbers.

A fundamental difference between these models is easily grasped for spatial structure in which human density differs among patches. In the density-dependent case, if mosquito density is the same across patches, each human is bitten at the same rate regardless of patch. In the frequency-dependent case, again for constant mosquito densities, humans are bitten at a lower rate in the larger patch (i.e., the patch with more humans); in the extreme, a parasite might be maintained only in the smaller patch because the biting rate per person is too low in the large patch. Only by scaling mosquito density with human density is it possible to maintain similar biting rates per humans across patches of different sizes.

Other models of biting dynamics have been developed, such as hybrid models that allow biting dynamics to vary across different host densities (Gandon, 2018; Xue et al., 2018). These models usually build in density dependence at one extreme of human density and frequency dependence at the other. Our use of models at both extremes obviates the need for hybrid models, given that we can show a fundamental difference between the two processes. The contrast of our models highlights the need to understand mosquito dynamics before reaching any conclusions about the impact of travel, and without an empirical basis for justifying even the hybrid models, hybrid models cannot be justified biologically any more than can the extremes.

### 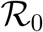 calculations when transmission is blocked in one patch

With vectored diseases, there are various ways to calculate the basic reproductive number, 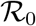 (e.g., Keeling and Rohani, 2008; Anzo-Hernández et al., 2019). Our method (Appendix) is essentially that of Keeling and Rohani (2008). For our purposes, the actual value of 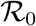 is unimportant, as we are interested in the relative impact on 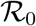 of changes in population structure, as well as a relative comparison of 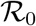 for density dependence and frequency dependence. Typically, different methods of computing basic reproduction numbers in vector models can lead to different 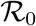 values (e.g., one value being the square of what is obtained via a different method) but they agree at the epidemic threshold of 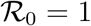, which again is the critical value between eradication and endemism.

To keep the focus on biological relevance, we limit consideration to 2 patches. As per our biological justification above, we let the intervention be fully effective and block all transmission in patch 1, but absent in patch 2. Maintenance of the parasite 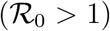 in this setting is due entirely to whether the parasite persists in patch 2.

We wish to consider conditions whereby, in the absence of human movement, the parasite would persist in patch 2. Our previous analysis (which neglected hosts and vectors, Bull et al., 2019) can be construed to suggest that, if patch 1 was sufficiently large, human movement between patches would facilitate parasite eradication by increasingly exposing the parasite to the average of both patches (as also true of Prosper et al. (2012)). We are interested in whether this conclusion holds: how does human movement affect persistence and how do the two models compare?

The 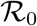 formula for either model (density-dependent or frequency-dependent) is a function of 7 parameters and 3 state variables (derived for general transmission values in the Appendix). The analysis assumes a small number of infected mosquitoes, an absence of infected humans in either patch, and no mosquito-to-human transmission in patch 1. For the frequency-dependent model with no mosquito-to-human transmission in patch 1, the formula is

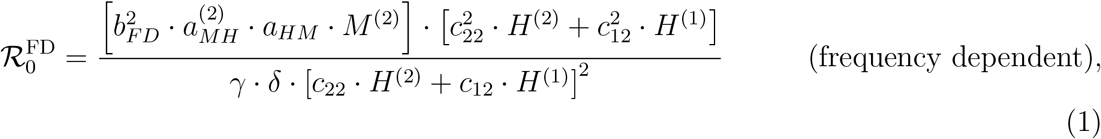

with notation defined in Table 1. The first numerator term in brackets is a mosquito term that accounts for the number of mosquitoes, transmission rates per bite in both directions, and biting rates; the squared biting rate accounts for the mosquito acquisition of the parasite and then its later transmission. The second numerator term in brackets is one of human population size weighted by (squared) human travel probabilities to account for only those humans present in patch 2—the patch with no block to transmission. The denominator is a squared term of humans present in patch 2, necessarily larger than the human term in the numerator given moderate or higher human densities (note that the *c*_*ij*_ ≤ 1). Inspection of this result reveals how increasing the numbers of humans in patch 2, while holding the mosquito term constant, reduces 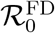, reflecting the dilution of mosquito bites. These results have been confirmed with limited numerical analyses of the full equations by varying the *c*_*ij*_. The threshold 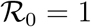 in (1) and (2) coincided with the threshold for maintenance or loss of the parasite.

**Table 1:** Description of state variables and parameters in the mathematical models.

| Notation | Description | Units |
| --- | --- | --- |
| $H^{(k)}$ | number of humans in patch $k$ | individuals |
| $M^{(k)}$ | number of mosquitoes in patch $k$ | individuals |
| $\gamma$ | recovery rate of infected humans | day<sup>-1</sup> |
| $b_{DD}$ | density-dependent biting rate | individual<sup>-1</sup>day<sup>-1</sup> |
| $b_{FD}$ | frequency-dependent biting rate | day<sup>-1</sup> |
| $\delta$ | mosquito death rate | day<sup>-1</sup> |
| $c_{kj}$ | fraction of time patch $k$ humans spend in patch $j$ | dimensionless |
| $a_{HM}$ | human to mosquito transmission probability | dimensionless |
| $a_{MH}^{(k)}$ | patch $k$ mosquito to human transmission probability | dimensionless |

What is of greater interest here is the comparison of 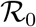 values between the density-dependent and frequency-dependent models. For the density-dependent model,

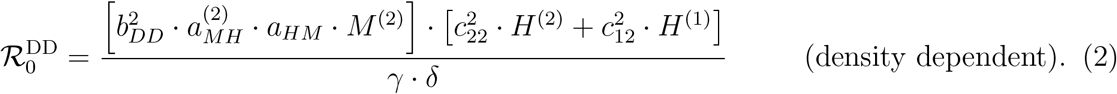

Note that the mosquito biting rate term here has different units than in (1)—see Table 1. Also note that there is no denominator term involving humans.

### Travel has different effects under frequency dependence versus density dependence

There are obvious similarities in the 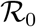 formulae, and we may compare them as follows:

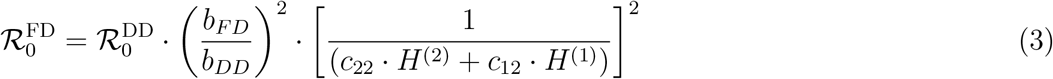

The difference of greatest biological interest is in the rightmost term of (3) when considered along with (1) and (2). With increasing numbers of humans in patch 2 (while maintaining constant mosquito density), the 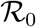 for frequency dependence declines, whereas the 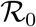 for density dependence increases. Note that increasing the number of humans in patch 2 can be accomplished either by increasing *c*_22_ or by increasing *c*_12_. Increasing *c*_22_ increases spatial structure globally, whereas increasing *c*_12_ reduces spatial structure.

For the goal of parasite eradication, which in both models requires its eradication in patch 2, the contrast between the frequency- and density-dependent models is extreme when considering spatial structure of humans. Reducing travel out of patch 2 increases 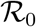 in the density-dependent case but decreases 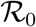 in the frequency-dependent case. The converse is true of travel into patch 2 (*c*_12_). Thus not only do the frequency-dependent and density-dependent models differ in the effect of changes in effective patch size (number of humans), but the effect of changing spatial structure differs between the models depending on whether travel involves humans leaving patch 2 or coming into it.

## Discussion

Our study is motivated by new technologies that are being used or will likely be used as interventions against vectored infectious diseases. They involve genetically modifying the vector to block its competence for parasite reproduction or transmission. As it is unlikely that any such interventions will cover entire vector populations, our interest lies in the consequences of unprotected vectors. In the absence of spatial structure, the overwhelming abundance of modified vectors would suppress the parasite, but with strong spatial structure, unprotected pockets/patches of vectors will enable the parasite to persist. What, then, is the effect of limited disruption of that spatial structure, as in the form of human travel?

We studied two types of well-established mathematical models of host-vector parasite dynamics. One model is a form of the long-used Ross-Macdonald model (e.g., McCallum et al., 2001; Keeling and Rohani, 2008; Prosper et al., 2012; Ruktanonchai et al., 2016; Soriano-Paños et al., 2020), a model that assumes frequency-dependent behavior of mosquito biting. Frequency dependence is characterized by individual mosquitoes biting at a fixed rate, less per person as the local human population increases. The other model used here is similar except in assuming density-dependent biting rates; here individual humans are bitten at the same rate per mosquito regardless of how many people there are. All models assumed 2 patches of humans and their resident mosquitoes; the mosquitoes in one patch were blocked from transmission, but the mosquitoes in the other patch were fully competent. With strict spatial structure (no human movement), the parasite would be completely absent in one patch but present at high levels in the other patch.

From a casual consideration of previous work (e.g., Prosper et al., 2012; Bull et al., 2019), we expected that any relaxation of spatial structure would reduce the disease 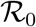 if the disease-free patch was large enough relative to the diseased patch. Thus sufficient human movement between the patches would eventually cause parasite extinction. The results were unexpected in that (i) different directions of movement (travel into or out of the patch) had opposing effects in a model, and (ii) those opposing effects were reversed between the two types of model. In hindsight, differences between the two models are understandable by considering the effect of increasing human density in a patch. In the density-dependent model, an increase in humans in a patch results in more mosquito biting (per mosquito); in the frequency-dependent model, mosquitoes do not increase biting activity and added humans results in a ‘swamping’ effect where most humans are protected due to the presence of other humans. These contrasting effects are at least broadly compatible with prior analyses that discovered opposing effects of movement on 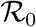 between frequency-dependent and density-dependent assumptions in single-population models (Wonham et al., 2006).

An obvious next step is to understand mosquito biting dynamics as it bears on disease transmission and human population structure. Simple extensions of frequency-dependent and density-dependent models may accommodate both behaviors as extremes in different biological realms, with high mosquito densities relative to humans tending toward density dependence, low densities tending toward frequency dependence (Gandon, 2018). However, additional complexities are possible: density and frequency dependence may differentially accrue to humans and vectors, and indeed, those two processes are not the only possible options for transmission dynamics (McCallum et al., 2001; Wonham et al., 2006).

Joint spatial structure of both vectors and humans is likely to present a major challenge to disease eradication by genetic modification of vector populations. Even with seemingly perfect blocking by the genetic engineering in those populations where it is implemented, potentially small unaltered vector populations will allow parasite maintenance provided the humans remain appropriately structured. Such pockets of escape may eventually be targeted for secondary interventions, but a major worry is that small pockets of persistence will become foci for evolution of resistant parasites that can then invade areas of more complete coverage. Understanding key dynamical properties of real systems may help predict which types of interventions can be combined to achieve local eradication.

## Appendix

### Two formulations of 2-patch vector-human models with cross-patch visits by humans: density-dependent vs. frequency-dependent transmission

Our models, written as systems of ordinary differential equations, track densities of susceptible and infected humans and mosquitoes in two patches connected by human movement. We let 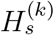 and 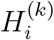 be the densities of susceptible and infected human hosts in patch *k* ∈ {1, 2} and 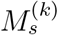 and 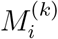 be the densities of susceptible and infected mosquitoes in patch *k*. The difference between the density-dependent and frequency-dependent models is encapsulated in the mosquito “biting rates”. For the density-dependent model, *b*_*DD*_ denotes the rate of biting per human per day by a given mosquito. In the frequency-dependent model, *b*_*FD*_*/H* denotes the rate of biting per human per day by a given mosquito when the (local) density of humans if *H*. In other words, a given mosquito doles out *b*_*DD*_ *H* bites per day in the density-dependent model, and *b*_*FD*_ bites per day in the frequency-dependent model. (When the number of humans increases, density-dependent mosquitoes work harder; frequency-dependent mosquitoes do not change their biting rate, but must allocate their bites among more humans.)

The probability that an uninfected human becomes infected when bitten by an infected mosquito from patch *k* is given by 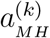. Dependence on the patch of the infecting mosquito reflects the assumptions that the level of parasite suppression is patch-dependent (as when the intervention is present in one patch but not the other) and an absence of mosquito movement among patches. Human-to-mosquito transmission is characterized by the parameter *a*_*HM*_, which denotes the probability that an uninfected mosquito becomes infected when it bites an infected human from either patch; there is no patch-specific interference of human-to-mosquito transmission. Said differently, patch-specific heterogeneity in transmission probability (and thus transmission rate) of the disease from mosquitoes to humans is what characterizes the effectiveness of the genetic intervention. Owing to the focus of intervention efforts on the transmission from vector to human host, there is no such need to introduce patch-specific differences in human-to-mosquito transmission.

Let *δ* denote the death rate of mosquitoes and *γ* the recovery rate of an infected human. We also let *λ_k_* be the birth rate of susceptible mosquitoes in patch *k*. The equilibrium density of mosquitoes in patch *k* is, thus, given by *λ_k_/δ*. We focus on parasite transmission dynamics when mosquito density is constant. The fraction of time a human residing in patch 1 spends in patch 1 (resp., patch 2) is denoted by *c*_11_ (resp., *c*_12_), where *c*_11_ + *c*_12_ = 1. Similarly, human residents of patch 2 spend fractions *c*_21_ and *c*_22_ in patches 1 and 2. Note that our human movement model is one of “visitation” rather than actual migration. An example would be people who commute between their home city and another for work. We assume that 0 ≤ *c*_12_ < 1 and 0 ≤ *c*_21_ < 1 to ensure that there are actually people in each patch.

### Density-dependent transmission

In the case of density-dependent transmission, infection rates for mosquitoes and humans have a mass-action dependence on mosquito and human densities.

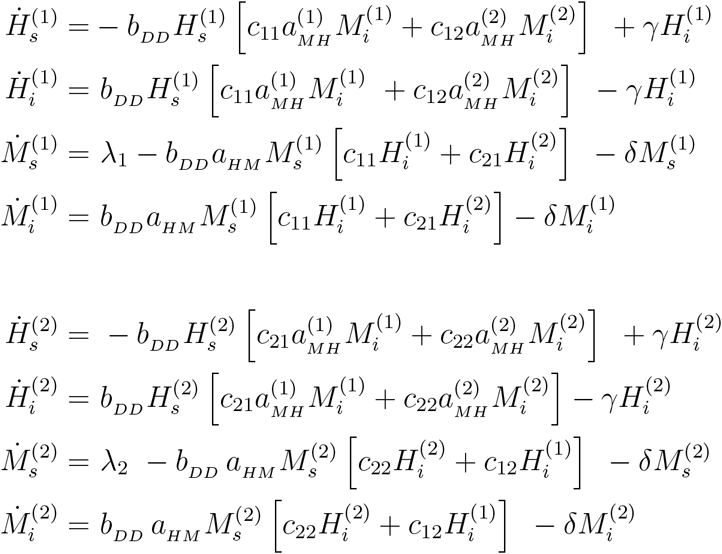

### Frequency-dependent transmission

Let 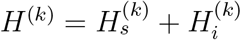 denote the total number of humans who reside in patch *k*. Then, for example, mosquitoes residing in patch 1 will see a mix of humans: *c*_11_*H*^(1)^ residents of patch 1 who are not visiting patch 2, and *c*_21_*H*^(2)^ residents of patch 2 who are visiting patch 1. The ‘effective’ number of humans in patch 1 (i.e., the number of humans experienced by mosquitoes in patch 1) is thus 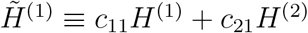. Similarly, the effective number of humans in patch 2 is 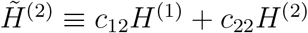. In the frequency-dependent transmission framework, a mosquito’s bites are randomly allocated to this mix of humans.

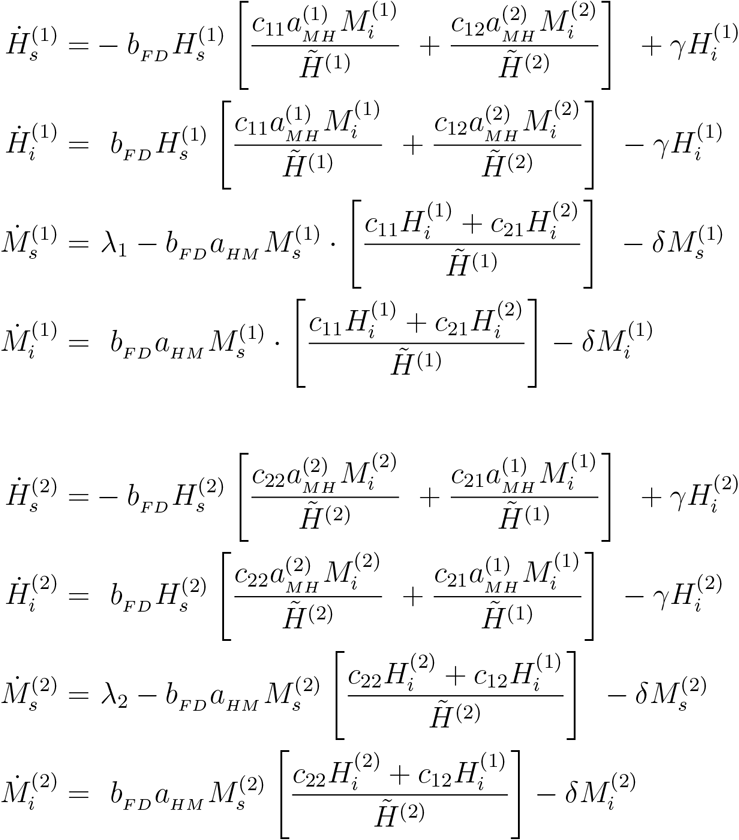

### Density-dependent 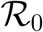 calculations

The basic reproduction number, especially for vectored disease models like those we consider, can be defined in several ways. These definitions give the same threshold condition 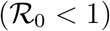 for the stability of the disease-free steady-state. Due to the multiphasic nature of vectored disease transmission, differences between definitions of the basic reproduction number for diseases like malaria can often be reconciled by realizing, say, one is the square of the other. A more fundamental issue in defining 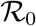 for mosquito-borne disease is the complexity that arises from having both human and vectors host the disease agent. Is 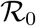 the number of secondary mosquito infections due to a small number of primarily infected mosquitoes in an otherwise susceptible population, or the number of secondary human infections due to a small number of initially infected humans, or some combination of the two? While the fates of mosquitoes and humans over the course of an epidemic are coupled, the mosquito- and human-centric basic reproduction numbers are indeed distinct quantities, agreeing only if the disease persists in the population at equilibrium.

To calculate the basic reproduction number 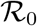 for the pathogen in mosquitoes, we assume that there is a small density of (primary) infected mosquitoes, 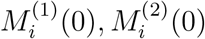, in patches 1 and 2, respectively, and no infected humans. In this initial phase, the density of susceptible mosquitoes is approximately *M*^(1)^ in patch 1 and *M*^(2)^ in patch 2, while the numbers of susceptible humans is *H*^(1)^ in patch 1 and *H*^(2)^ in patch 2. To compute the numbers of secondary infections of mosquitoes in each patch, we must consider two steps: mosquito-to-human followed by human-to-mosquito transmission.

1. The number of humans directly infected from primary mosquitoes before they die is:

- in patch 1:

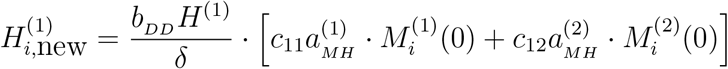
- in patch 2:

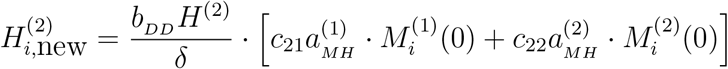
2. The number of mosquitoes infected by these newly infected humans before they recover is:

- in patch 1:

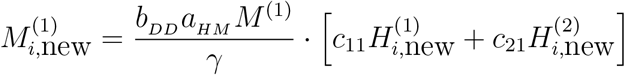
- in patch 2:

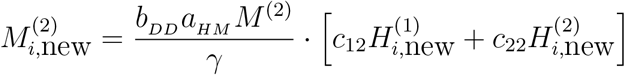

Note that mosquito death rate *δ* corresponds to mean lifetime 1*/δ*; similarly, 1*/γ* corresponds to the mean time before an infected human recovers. Combining the above two steps allows us to specify patterns of secondary infection (per primary infected mosquito in each patch) in the matrix

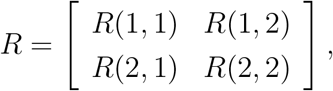

where *R*(*j, k*) denotes the number of secondary mosquito infections in patch *j* that arose from primarily infected mosquitoes in patch *k*, for *j, k* ∈ {1, 2}. Consequently, the *j*th row sum gives the number of secondary mosquito infections in patch *j*, and the *k*th column sum is the total number of secondary infections due to initially infected mosquitoes in patch *k*. Tracking the patterns of infection in both patches, we find that

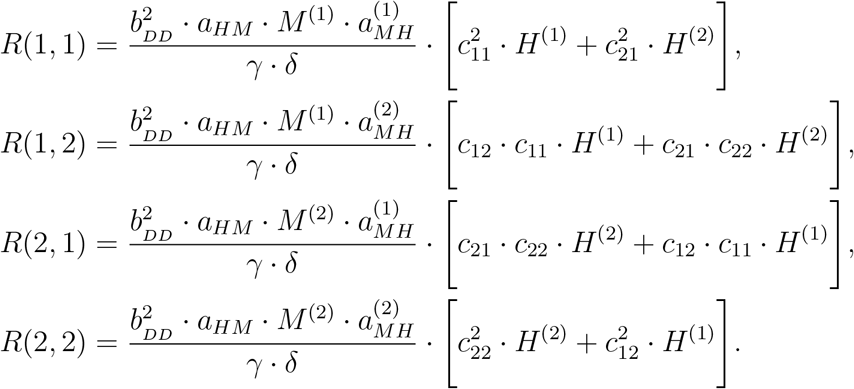

Note that each of the four secondary transmission terms above has two components: one corresponding to a susceptible human from patch 1 being infected by a primary infected mosquito from the designated patch, and one corresponding to a susceptible human from patch 2 being infected by a primary infected mosquito. Recall that mosquitoes are tied to their patch; only humans visit the other patch. For example, *R*(1, 2) records the number of secondary infections of mosquitoes living in patch 1 that arose from a primary infected mosquito in patch 2. There are two patterns of human visitation that can lead to this event. (1) encoded in the term *c*_12_*c*_11_*H*^(1)^ on the right-hand side of the *R*(1, 2) expression: in the first phase, a human in patch 1 visits patch 2 and is infected by a primary mosquito there (and the human returns to its home patch); in the second phase, the newly infected human stays in patch 1 and infects a susceptible mosquito there. (2) encoded in the term *c*_22_*c*_21_*H*^(2)^ on the right-hand side of the *R*(1, 2) expression: in the first phase, a human in patch 2 remains in patch 2 and is infected by a primary mosquito there; in the second phase, the newly infected human visits patch 1 and infects a susceptible mosquito there.

The *basic reproduction number* 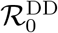 *for the density-dependent transmission model* is the leading eigenvalue of the matrix *R*. The special case of no mosquito-to-human transmission in patch 1 (i.e.,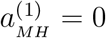) is interesting in that *R*(1, 1) = 0 = *R*(2, 1) and hence 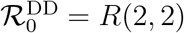.

### Frequency-dependent 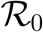 calculations

Similar to the above case, the calculation of a mosquito-centric 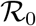 for the frequency-dependent case begins with an assumption that there is a small density of (primary) infected mosquitoes, 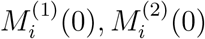, in patches 1 and 2, respectively, and infected humans. In this initial phase, the density of susceptible mosquitoes is approximately *M* ^(1)^ in patch 1 and *M* ^(2)^ in patch 2, while the numbers of susceptible humans is *H*^(1)^ in patch 1 and *H*^(2)^ in patch 2. To compute the numbers of secondary infections of mosquitoes in each patch, we must consider two steps: mosquito-to-human followed by human-to-mosquito transmission.

1. The number of humans directly infected from primary mosquito before it dies is:

- in patch 1:

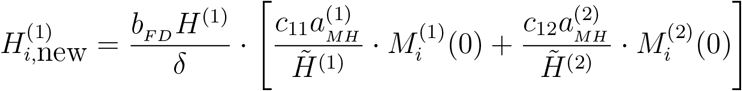
- in patch 2:

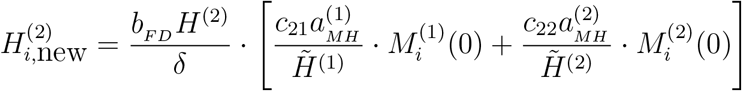
2. The number of mosquitoes infected by these newly infected humans before they recover is:

- in patch 1:

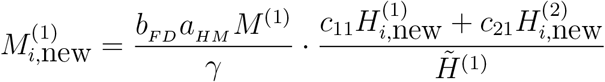
- in patch 2:

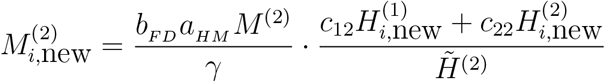

Putting these together allows us to specify patterns of secondary infection (per primary infected mosquito in each patch) in the matrix

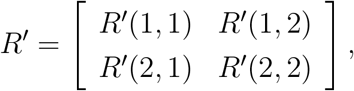

where *R*(*j, k*) denotes the number of secondary mosquito infections in patch *j* that arose from primary infected mosquitoes in patch *k*, for *j, k* ∈ {1, 2}. Tracking the patterns of infection in both patches, we arrive at

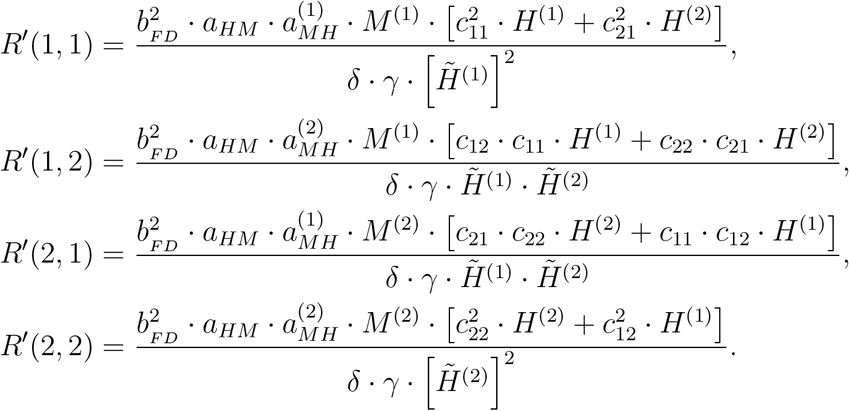

The *basic reproduction number* 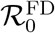 *for the frequency-dependent transmission model* is the leading eigenvalue of the matrix *R*^′^. As in the density-dependent transmission model, the special case of no mosquito-to-human transmission in patch 1 (i.e., 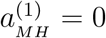 results in *R*^′^(1, 1) = 0 = *R*^′^(2, 1) and hence 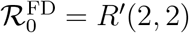. If we assume both 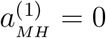 and *c*_12_ = 0, then we obtain a stark difference between these models: 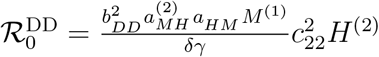 for the density-dependent model, and 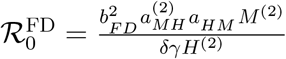 for the frequency-dependent model. Thus, one-way visitation to a patch with perfect cargo has a strong effect in the density-dependent model, but no effect in the frequency-dependent model. In fact, the latter 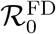 is in the standard form for a Ross-Macdonald model with no patch structure.

Notice that mosquito and human densities in the terms characterizing the basic reproduction number in the frequency-dependent model appear in ratio form *M/H*, while in the density-dependent model they appear in product form *MH*.

Our 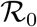 calculations, for both density- and frequency-dependent transmission, were based on computing numbers of secondarily infected mosquitoes that arose from the primary mosquito infections. Since human and mosquito infections are intertwined due to the nature of vector transmission, it should not be surprising that the threshold 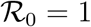 above which human infection persists is the same as the one that guarantees persistence of mosquito infection. In numerical solutions of our differential equations (not shown), we saw positive equilibrium densities of both infected mosquitoes and infected humans precisely when 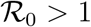.

## Notes

### Competing Interest Statement

The authors have declared no competing interest.

